# A rapid urban biodiversity blitz using aquatic environmental DNA

**DOI:** 10.1101/2020.05.27.116905

**Authors:** Kamil Hupało, Markus Majaneva, Molly Victoria Czachur, Lucas Sire, Daniel Marquina, Darío A. Lijtmaer, Vladislav Ivanov, Sonja Leidenberger, Fedor Čiampor, Zuzana Čiamporová-Zaťovičová, Izabela S. Mendes, Andrea Desiderato, Lasse Topstad, Kenny Meganck, Danial Hariz Z. A., Gaute Kjærstad, Xiao-Long Lin, Benjamin Price, Mark Stevens, Torbjørn Ekrem, Kristy Deiner

## Abstract

**Background:** As global biodiversity declines, there’s an increasing need to create an educated and engaged society. Having people from all ages participate in measuring biodiversity where they live helps to create awareness. Recently, the use of environmental DNA (eDNA) for biodiversity surveys has gained momentum. Here, we test whether sampling eDNA and metabarcoding can be used for rapid urban biodiversity surveys for educational purposes.

**Materials & Methods:** We sampled 2×1 L of water from each of 15 locations in the city of Trondheim, Norway, including a variety of freshwater, marine and brackish habitats. DNA was extracted, amplified in triplicate for the COI gene and sequenced. The obtained data were analysed on the novel mBRAVE platform, an online open access software and computing resource.

**Results:** The water samples were collected in two days by two people and the lab analysis was completed in five days by one person. Overall, we detected the presence of 501 taxa identified as belonging to 435 species, representing 90 orders and 18 phyla. On average, only 5.4% of the taxa were shared among six replicates per site. Based on the observed diversity, three distinct clusters were detected and related to geographic distribution of sites. There were some taxa shared between the habitats, with a substantial presence of terrestrial biota.

**Discussion:** Our results match expected patterns of biodiversity in the landscape and show that with minimal sampling effort, hundreds of species can be detected. Thus, using eDNA analysis of water is promising for rapid biodiversity surveys, and it is likely that more detailed results could be obtained by optimising field and lab methods for particular groups of interest. We recommend that rapid eDNA surveys, with openly available services and softwares, can be used to raise awareness in the importance of biodiversity.

## 1 Introduction

Global biodiversity is imperiled with an estimated one million species at risk of extinction (IPBES, 2019). There is an urgent need to build consensus around action and reverse the impending loss of life from our planet. Conducting rapid biodiversity assessments in urban centres such as the City Nature Challenge (http://citynaturechallenge.org/) are increasingly used as means by which to educate all ages and sectors of society about the importance of biodiversity. Many biodiversity observation programs exist to engage with the public (e.g., Bio-Blitz; Postles & Bartlett, 2018). However, when such surveys occur, the amount of equipment, resources and time needed from experts to survey diversity across the tree of life quickly becomes intractable for many educators. Thus, educational and outreach programs using rapid biodiversity surveys are often limited in their taxonomic breadth, offered mostly by larger organizations, are infrequent, and their success depends largely on the engagement of the organizers (Ruch et al., 2010; Postles & Bartlett, 2018).

New DNA-based technology has the power to solve current limitations of education programs wanting to teach about biodiversity through first hand experiences. Namely, in the last decade, the scientific community has realized we can survey species by finding trace amounts of their DNA left behind in the environment (i.e., environmental DNA ‘eDNA’) and have gained tremendous momentum (e.g., Deiner et al., 2017; Cristescu & Hebert, 2018; Thomsen & Sigsgaard, 2019). If biodiversity surveys using DNA from the environment are easy to apply, then they can be used as an educational outreach tool for engaging the public with first hand biodiversity knowledge to create awareness. Furthermore, because environmental DNA sequencing services are common and commercially available from many companies around the world, access for educators at all levels is possible.

To test whether eDNA surveys can be used in an urban setting for educational purposes, we performed a test case in the city of Trondheim, Norway (Figure 1). Trondheim is situated at the Trondheim Fjord in Central Norway at 63° North. It has almost 200,000 inhabitants and is the third largest city in Norway. Although the city centre is rather developed, there are numerous parks mixed within residential areas, and outside of the city centre itself there are gardens and small natural forest patches. To the west, the city is bordered by the forest Bymarka, partially a nature reserve; while to the east, the city is bordered by farm fields and the forest Estendstadmarka. Both these natural areas have a rolling landscape, with the highest peak Storheia at 565 m above sea level. They contain the headwaters of multiple streams running towards the city centre and the Trondheim Fjord. The river Nidelva originates from the large lake Selbusjøen to the south-east, splits the city and empties into the fjord north of Trondheim. The river is about 31 km long, is regulated for hydropower purposes, and the water discharge level is heavily influenced by frequent variation in electricity production intensity (‘hydropeaking’). However, a minimum discharge of 30-40 m^3^/s is required, partly to ensure that the viable trout and salmon populations in the river have acceptable living conditions. The river, streams and ponds in Trondheim are influenced by human activity to a variable degree. In some areas, the aquatic habitats are strongly influenced by land use (e.g., road infrastructure), while other sections are influenced by sewage overflow in the drainage system. In 2016, the city of Trondheim treated seven lakes in Bymarka with rotenone to remove the invasive common roach (*Rutilus rutilus* (Linnaeus)). Several ponds around Trondheim are known to host great-crested newt (*Triturus cristatus* (Laurenti)) and common newt (*Lissotriton vulgaris* (Linnaeus)), but their presence near the city centre has declined due to habitat alteration (Tilseth, 2008). The Trondheim Fjord was heavily polluted by heavy metals and organic pollutants but has improved considerably after a large-scale clean-up and dredging in 2015-2016.

**Figure 1:**
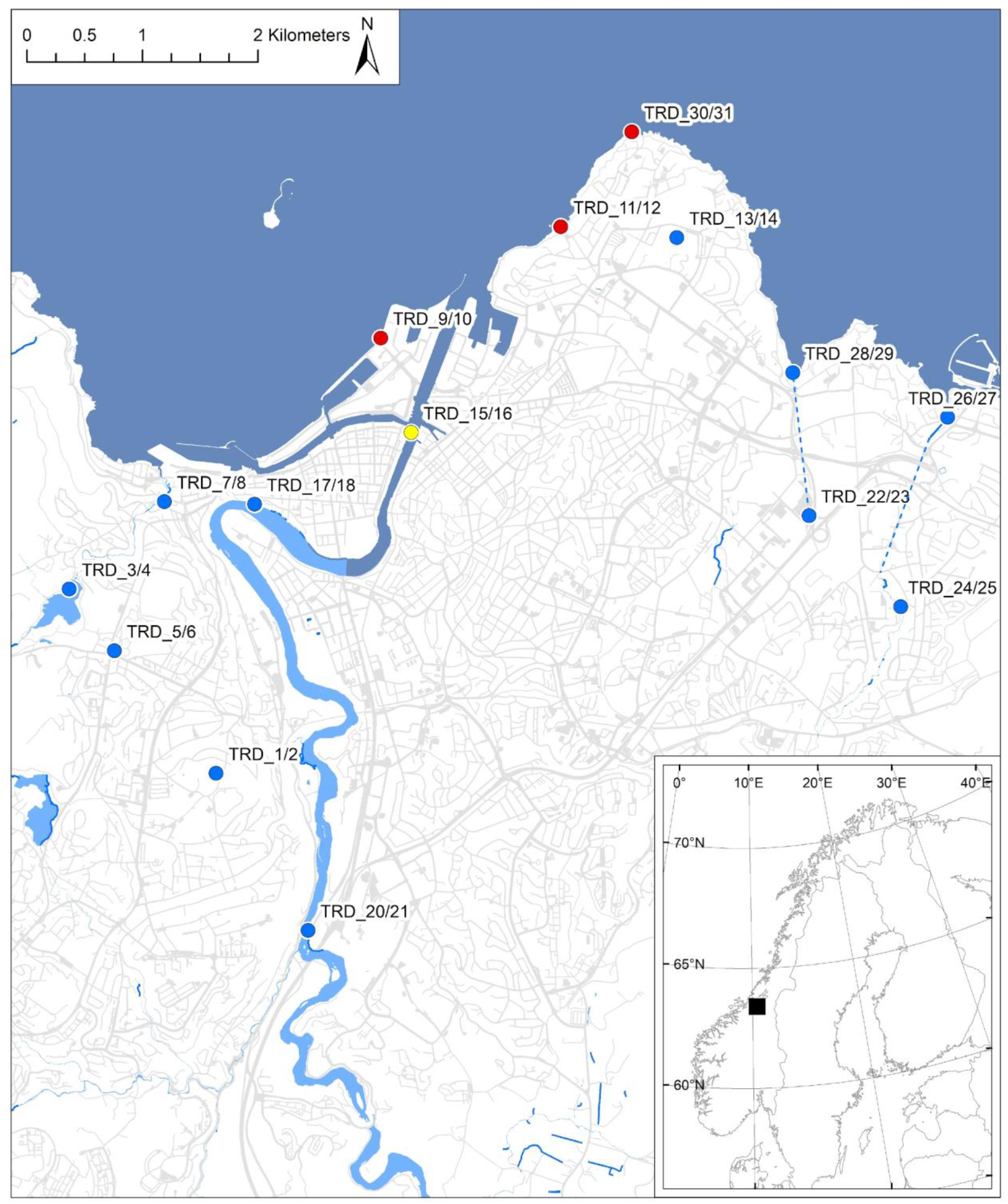
The 15 sampled locations surveyed around the urban centre of Trondheim on May 13 and 14, 2019 [TRD_1/2: pond Havstein golf course; 3/4: Theisendammen; 5/6: pond Sverresborg Museum; 7/8: Ilabekken at Bleikvollen; 9/10: Pirsenteret; 11/12: Korsvika; 13/14: Ringve Botanical Garden; 15/16: Nidelva at Trondheim Maritime Museum (downstream); 17/18: Nidelva at Fylkesmannsboligen (midstream); 20/21: Nidelva at Sluppen (upstream); 22/23: Madsjøen at IKEA; 24/25: Stokkbekken upstream; 26/27: Stokkbekken downsteam; 28/29: Leangenbekken; 30/31: Østmarkneset at Ladekaia; 19,32: blank, thus excluded from the map]. The color of the dots corresponds to the sites’ habitat: blue - freshwater sites, red - marine, yellow - brackish. The dotted lines represent the underground connections between the sites.

With this backdrop, we tested whether eDNA could be rapidly sampled and sequenced from 15 sites and taxa lists determined through use of DNA-based identifications of observed sequences to the BOLD database (Ratnasingham & Hebert, 2007) using the novel Multiplex Barcode Research And Visualization Environment (mBRAVE) platform (http://www.mbrave.net/). To make use of the data and show its application for educational purposes, we used our rapid eDNA biodiversity assessment method to determine patterns in observed biodiversity. We asked i) how much total diversity could be detected at the selected locations and, ii) whether patterns of diversity changed due to sampling location or habitat. Additionally, we wanted to iii) test whether we could detect rare amphibians: the great-crested newt (*Triturus cristatus*) and the common newt (*Lissotriton vulgaris*), and iv) an invasive fish: the common roach (*Rutilus rutilus*). Lastly, v) we asked whether eDNA was transported across the landscape and between habitats.

## 2 Materials & Methods

### 2.1 Sampling

Two 1 L water samples were collected from each of 15 sites in Trondheim, Norway (Figure 1) on May 13^th^ and 14^th^, 2019 (Table S1). The sites included ponds, small streams, a lake, rivers, and the sea. At each site, water was collected by submerging two sterilized (5% bleach, flushed with the sampling site water) 1 L rectangular polyethylene terephthalate bottles (Nalgene/VWR International, Radnor, PA, USA) just below the surface, with only bottle and gloved hand touching the water. At the running water sites, water was collected upstream of where the collector was standing. Samples were stored in a cooler box. The air temperature was between 0-10 °C, so the samples were kept at approximately ambient temperature until filtered.

### 2.2 Clean Laboratory environment

The samples were processed in a unidirectional controlled facility. Filtration was done in a separate room, where no other environmental or organismal samples were processed, and all surfaces were decontaminated before use (with a 5% bleach solution and 70% ethanol). Each filter was folded with a fresh pair of forceps, which were decontaminated overnight in 5% bleach solution before the second day of filtration. The DNA extraction was done in a laboratory where all surfaces and all necessary equipment were decontaminated with a 5% bleach solution and 70% ethanol, while wearing a full-body suit, face-mask and double gloves. After a workday, UV-light was set to run overnight. Respective DNA extracts were transferred to a pre-PCR room where PCRs were set up. After amplification, the libraries were prepared in the pre-PCR room and the template was added in the post-PCR room.

### 2.3 Sample filtration and DNA extraction

Samples (N = 30) were transferred to the local university laboratory (NTNU University Museum, Norwegian University of Science and Technology) within two hours after sampling. All samples were filtered on the same day they were collected and in the order they were sampled using an electrical vacuum pump connected to a manifold (Pall Laboratory, Port Washington, NY, USA) carrying three individually operated filter holder bases. Samples were filtered through sterile and individually packed 0.45 μm mixed cellulose ester filters (ø 47 mm) attached to a 300 mL reservoir (Pall Laboratory). Filters were folded inward three times with decontaminated forceps and placed into 1.5 mL centrifuge tubes with 800 μL of ATL lysis buffer (Qiagen GmbH, Hilden, Germany). To control for contamination during the filtration process on each day a negative control was made by filtering 500 mL of molecular grade water in the same manner as samples after all field samples were processed (Table S1). Hereafter these are referred to as sample blanks. Prior to DNA extraction, the lysis buffer was removed from the filter and partitioned into two tubes, each containing ~400 μL of buffer ATL. The DNeasy Blood and Tissue kit (Qiagen) was used for DNA extraction from the lysate according to the manufacturer’s instructions for tissues with the exception that they were shaken horizontally at 250 rpm at 37 °C overnight. The purified DNA was eluted with 200 μL of molecular grade water. No negative DNA extraction controls were created, but the sample blanks served as a full process controls for all laboratory procedures. Quantification of the DNA extracts was not performed.

### 2.4 PCR amplification and sequencing

The extracted DNA from each of the two samples from the 15 sites and two sample blanks (N = 32) were amplified using a common two-step PCR method, with the highly degenerate BF2-BR2 primers (Elbrecht & Leese, 2017) with attached Illumina (San Diego, CA, USA) adapters 5’-TCGTCGGCAGCGTCAGATGTGTATAAGAGACAG-3’ (forward) and 5’-GTCTCGTGGGCTCGGAGATGTGTATAAGAGACAG-3’ (reverse) in the first PCR. The primers target an approximately 420 bp long fragment of the mitochondrial cytochrome *c* oxidase subunit I gene (COI). Each DNA sample was amplified in three technical PCR replicates and a negative control was created using molecular grade water in place of the template. Each PCR had a final volume of 25 μL containing 2 μL DNA template (DNA was diluted 1:10 prior to amplification to reduce the chance of inhibition), 17.8 μL molecular biology grade water, 2.5 μL 10× reaction buffer (200 mM Tris HCl, 500 mM KCl, pH 8.4), 1 μL MgCl2 (50 mM), 0.5 μL dNTPs mix (10 mM), 0.5 μL of each primer (10 mM), and 0.2 μL Platinum Taq polymerase (5 U/μL) (Invitrogen). The PCR conditions were, with a heated lid, 94 °C for 5 min, followed by a total of 35 cycles of 94 °C for 40 s, 50 °C for 1 min, and 72 °C for 30 s, and a final extension at 72 °C for 2 min. PCR products were visualized on a 1.5% agarose gel to check the amplification success.

In the second PCR, the Illumina tailed amplicons (N = 96) were dual indexed (Supplementary File 1), using Nextera XT Index 1 and 2 primers and Nextera XT Index v2 set D (Illumina, FC-131-2004) in a reduced-cycle PCR (15 cycles) according to the manufacturer’s protocol. The amplification reactions contained the same reagent concentrations as above with three modifications: 3 μL of product from the first PCR, 1 μL of each primer and 0.25 μL of Platinum Taq (Invitrogen). PCRs were not purified between the first and second amplification. Indexed amplicons were pooled by equal volume (5 μL each) into a single library. The library was purified and size-selected (>200 bp fragments retained), using SPRIselect (Beckman Coulter) with a ratio of 0.92. Library was quantified using Qubit™ dsDNA HS Assay Kit (ThermoFisher), following the manufacturer’s protocol. Library was loaded at a 14 pM concentration with 6 % PhiX added. Sequencing was performed on a MiSeq v3 flow cell sequencing paired end 2×300 bp and index 8 + 8 bp at the Genomics Core Facility, Oslo University Hospital, Oslo, Norway. The run folder was imported into the Illumina Local Run Manager version 2.0.0 and was used to complete the base calling and demultiplex the libraries to generate the fastq files for each sample (N = 96).

### 2.5 Data upload, filtering criteria and taxonomic assignment

The resulting 96 fastq files from the Illumina MiSeq run were imported into mBRAVE using the sample batch function. Each demultiplexed library (i.e., fastq file) is now referred to as a ‘TRD run’ for downstream analysis. Every parameter described here was retrieved from the mBRAVE platform and was available to the user as last accessed in July 2019. An extended description of the methods can be accessed in Supplemental File 1. Thus, we briefly describe the methods used to format, clean, and denoise the raw sequence data before taxonomic identification and biodiversity analyses.

For each library now consisting of a TRD run, the paired end merging of MiSeq reads required a minimum 20 bp overlap between the forward and reverse reads, while allowing up to 5 nucleotide substitutions. Both front and end part of the resulting sequences were trimmed for 20 bp to remove the primer sequence, followed by a trimming of total sequence length down to 500 bp. Low quality sequences were removed if the average quality value (QV) was less than 20 or sequences were shorter than 200 bp. This filtering step allowed for a max of 2% nucleotides with >20 QV value and max 1% nucleotides with >10 QV value. Sequences fulfilling these criteria were de-replicated and clustered on mBRAVE as Operational Taxonomic Units (OTUs) using a 2.5% similarity threshold. Obtained OTUs were taxonomically assigned using an initial 2% ID distance threshold to the reference sequences. When the obtained sequences could not be assigned to any reference sequence, the threshold levels were relaxed until a hit was found for higher taxonomic ranks: genus (3%), subfamily (4%), family (5%), order (7%), class (10%) and phylum (12%). The publicly available BOLD reference libraries for Insecta, non-insect

Arthropods, non-arthropod invertebrates, Chordata and bacteria, as well as a standard contamination reference database, were used for identification of all OTUs. Whenever a taxonomic assignment from a reference sequence returned more than one species name, taxonomy was assigned to higher taxonomic ranks with no conflicting taxonomy.

After selecting parameter values, mBRAVE automatically applied the same parameters to each TRD run in our dataset. For each run, a summary was generated and checked to ensure good fit with our filtering criteria. Data visualisations within the mBRAVE environment allowed for the data to initially be reviewed. These visualizations included sequence length distribution, GC composition distribution, run QV score distribution and BIN (Barcode Index Numbers, Ratnasingham and Hebert, 2013) count vs OTU count. Where low quality sets were observed, the low-quality runs were removed. The parameters chosen for this study were adjusted after considering the quality comparison in mBRAVE. Data were then combined into ‘sets’ within the mBRAVE environment, where biological and technical PCR replicates for each sample site were combined as a single set that could be further analysed.

### 2.6 Biodiversity and distribution analyses

To assess general patterns of diversity across the city of Trondheim, a summary report for all sets was generated to calculate the number and list of taxa detected in each TRD run. In order to have an overview of which taxa (i.e. BINs) were shared among samples and the negative filter controls, the beta diversity tool with the Jaccard similarity index was used within the mBRAVE environment using the sample sets as units of comparison. A heatmap was generated with either the shared BINs or the index value between the paired sets.

To document the composition of observed diversity in each run, it was considered that freshwater, marine, brackish and terrestrial taxa could be observed. Thus, sites were designated as belonging to one of three different realms (marine, freshwater, brackish) (Table S1). A brackish site was designated as such due to the tidal activity and temporal admixture of freshwater and seawater. When taxa could be assigned to a species, the known habitat designation was indicated, which included marine, freshwater, brackish and terrestrial using the WoRMS database (World Register of Marine Species, Horton et al., 2017) or other internet sources and in some cases, the species’ original descriptions. When taxa could not be assigned to species, the sequences and records were cross-validated with the following public databases: GBIF (https://www.gbif.org/); Fauna Europaea (https://fauna-eu.org/) and NCBI GenBank (https://www.ncbi.nlm.nih.gov/genbank/)

To assess similarity within sites among the biological and technical replicates, diversity at a local scale was evaluated with the BIN comparison tool in mBRAVE. This generated a Venn diagram with unique and shared BINs. Two approaches were adopted, the first without excluding any BIN, and secondly by accepting only the BINs shared at least by two replicates (biological or technical). We present only the data based on the latter.

Beta diversity analysis was performed in mBRAVE for all TRD runs using the beta diversity grouping runs by sets, composed of all replicates belonging to a particular site or habitat. To assess the impact of singleton/doubleton removal, the beta diversity analyses were run for every BIN (not excluding those found in less than two replicates), and this analysis was run again excluding singletons and doubletons (i.e., selecting in *Minimum Sequences per BIN or Species* option: 1, 2 or 3 respectively). Heatmaps and similarity measures were generated for the selected samples during the beta diversity analysis. We further explored the effect of including contamination BINs, as indicated by the presence of BINs in sample blanks runs, or the effect of BINs with unclear taxonomic resolution on the diversity observed for all sites.

## 3 Results

### 3.1 General patterns of diversity and similarity across the city of Trondheim

Overall, we detected the presence of 501 BINs identified as belonging to 435 species, representing 90 orders and 18 phyla; 9 BINs could not be assigned to any known taxa (Figure 2, Supplemental File 2). The full list of OTUs with their associated BINs, taxonomy and habitat associations are in Table S2. Many OTUs were obtained that could also not be associated with a BIN. These OTUs were not considered further in the analysis. We also note that not all data in the BOLD reference library are public; therefore, some taxonomic names associated with a particular BIN were unavailable when searched through BOLD itself and can only be accessed via mBRAVE. Most of the detected diversity consisted of Arthropoda (312 BINs), with Diptera being the most represented order (141 BINs). The least represented taxa (at most 2 BINs per taxon) were Actinobacteria, Bacteroidetes, Chaetognatha, Gastrotricha, Phoronida, Porifera and Tardigrada. The most common taxon was *Candidatus pelagibacter* Rappé, Connon, Vergin & Giovannoni, present in 12 out of 15 sites (Table S2).

**Figure 2:**
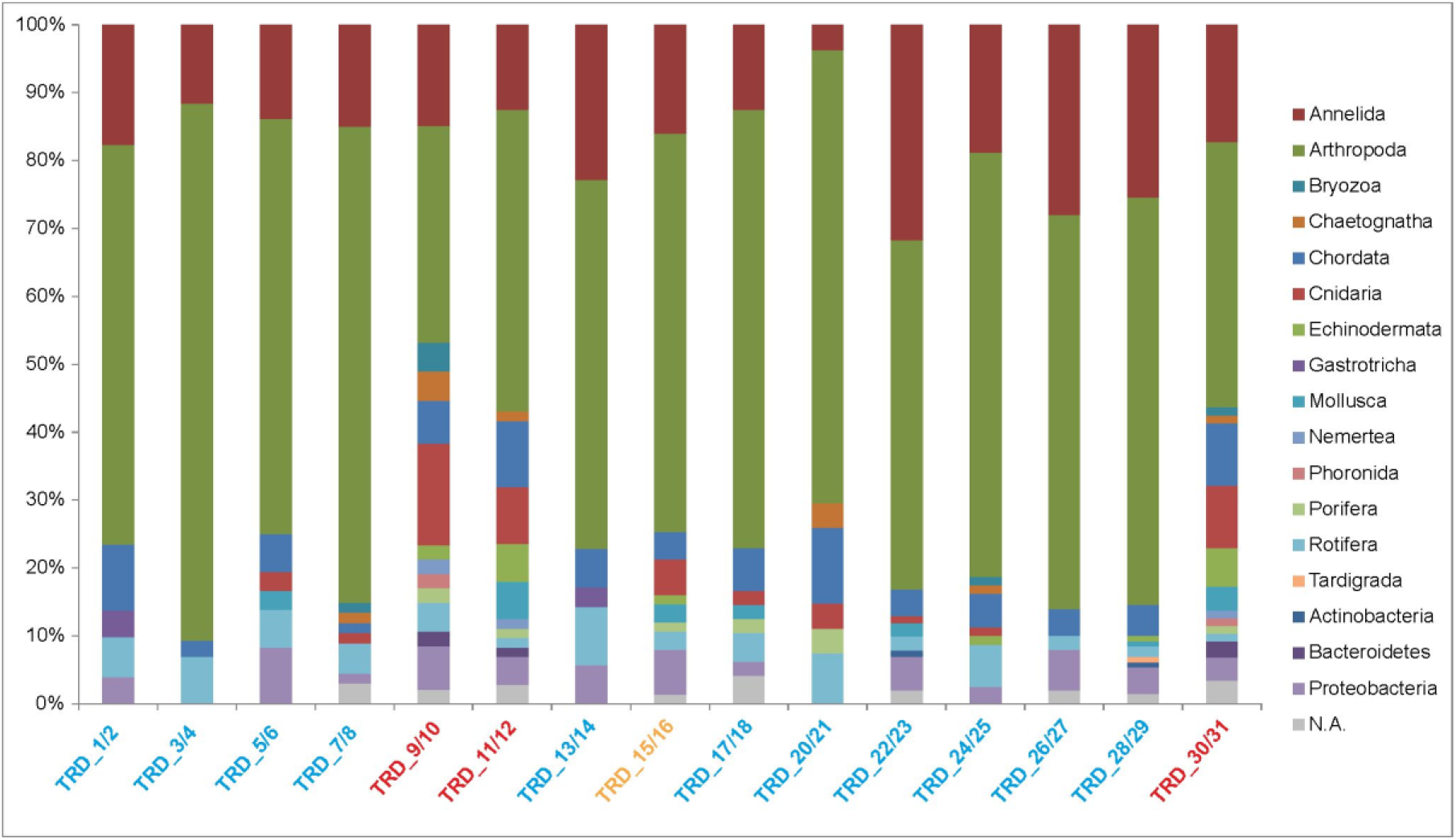
The normalized taxonomic composition of each studied site according to the Barcode Index Number (BIN) assignment done to phylum level. Colors of site names correspond to those used in Figure 1.

According to the Jaccard similarity index, the sites were clustered in three groups, Group A: pond Havstein golf course (TRD_1/2), ponds at Sverresborg Museum (TRD_5/6) and Ringve Botanical Garden (TRD_13/14); Group B: Theisendammen (TRD_3/4), Ilabekken at Bleikvollen (TRD_7/8), Madsjøen at IKEA (TRD_22/23), Stokkbekken upstream (TRD_24/25), Stokkbekken downstream (TRD_26/27) and Leangenbekken (TRD_28/29); and Group C: Pirsenteret (TRD_9/10), Korsvika (TRD_11/12), Nidelva at Trondheim Maritime Museum (downstream; TRD_15/16), Nidelva at Fylkesmannsboligen (midstream; TRD_17/18), Nidelva at Sluppen (upstream; TRD_20/21) and Østmarkneset at Ladekaia (TRD_30/31). Both Groups A and C formed one cluster each in the dendrogram based on the values of the Jaccard similarity index between samples (Figure 3), but four samples of Group B clustered together with Group C, separated from the other two samples of Group B. Group A was the most homogeneous and distinct group with Jaccard similarity indices that ranged from 0.12 to 0.16 among samples within group, and 0.009 to 0.093 among samples between groups, followed by Group C (with ranges of 0.096 – 0.23 and 0.23 – 0.15 within and between groups respectively), while in Group B (ranging 0.036 – 0.26 and 0.023 – 0.15), some samples showed greater similarity with samples from Group A than with other samples within Group B. Moreover, the sample blanks were different from the rest of the sites, forming a separate, fourth group. The highest number of BINs (130) were detected in Leangenbekken (TRD_28/29) with the lowest number of BINs (27) detected in the upstream site Sluppen of the Nidelva river (TRD_20/21) (Figure 3).

**Figure 3:**
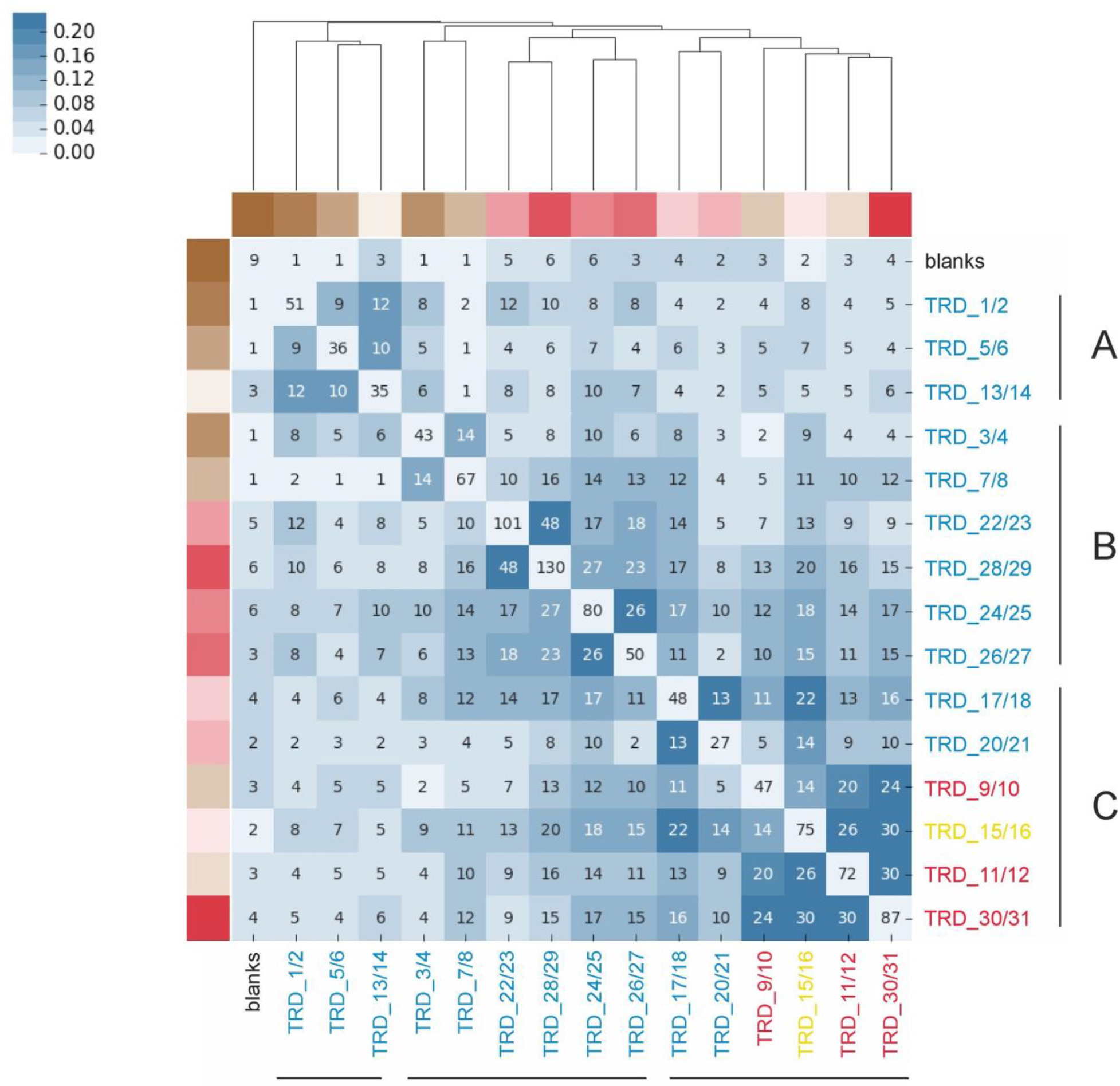
Heatmap generated in mBRAVE for Jaccard’s similarity index among samples where the darker the blue, the greater the similarity is between sampled sites. Clusters indicated with a dendrogram on top. Numbers correspond to the number of Barcode Index Numbers (BINs) identified within and among sites. The lines and letters correspond to the group clustering. Site names’ colors correspond to those presented in other figures.

### 3.2 Similarity within sites among biological and technical replicates

The similarity between biological and technical replicates for each site was low (Figure 4). On average, only 5.4% of the BINs were shared among all six replicates for each sampling site (2 biological replicates and 3 technical replicates for each of these). The highest percentage of shared BINs (Pirsenteret/TRD_9/10) was 15%, with two cases (Nidelva at Sluppen/TRD_20/21 and Stokkbekken downstream/TRD_26/27), that did not share BINs among replicates. The number of shared BINs was further impacted by whether BINs were represented by only one or two sequences in the full dataset. However, this mostly affected the sharing of BINs in the blanks with the rest of the sampled sites (Supplemental File 2).

**Figure 4:**
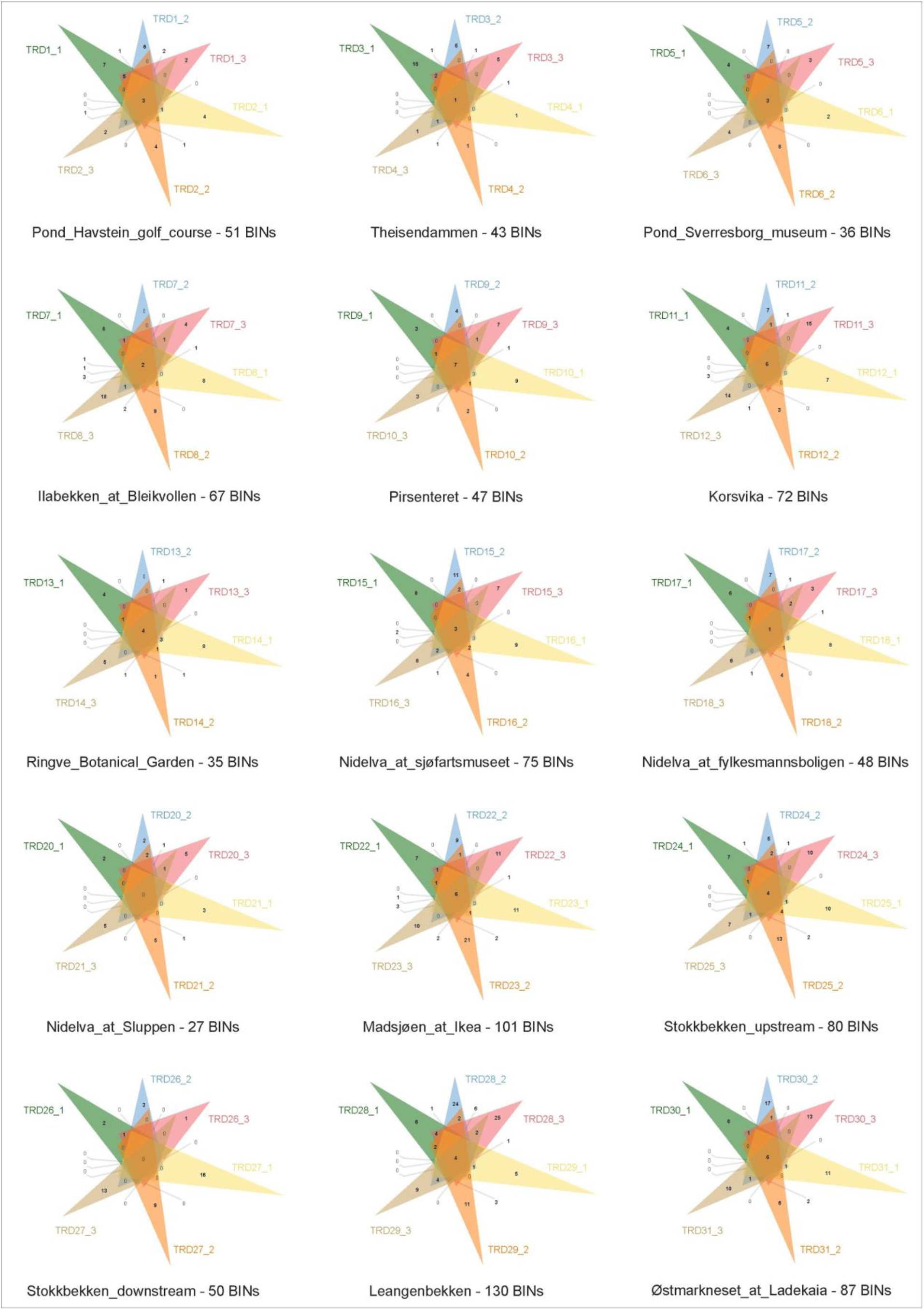
Venn diagrams generated in mBRAVE for each site showing the total number of Barcode Index Numbers (BINs) and number of BINs shared within sites among biological replicates (indicated with ‘TRD’ and a corresponding site name) and technical replicates (indicated by _1, _2 or _3).

Among the sample blanks, nine BINs were detected (Supplemental File 2). All of them were present in other samples (Table S2). The number of BINs observed and the number of reads for any BIN was low. When BINs represented by a single read (singleton) or two reads (doubletons) were to be removed, only one BIN remained. Thus, we did not consider this as evidence of extensive lab contamination during the processing of samples. Instead, it is more suggestive of sequencing errors in the indexes resulting in a small number of reads being misassigned to a sample when samples were demultiplexed in the bioinformatic processing (known as tag jumping). As this error is random and low, it is unlikely to have a major impact on our results of similarity among samples. Thus, no removal of BINs was done prior to further analysis. In addition to many BINs having few representative sequences, we observed taxonomic incongruences of species names with BINs in the BOLD reference database (presented in detail in Supplemental File 2 and Table S3).

### 3.3 The composition of observed diversity across sites - freshwater, marine, brackish and terrestrial taxa

On the 11 freshwater sites, 396 BINs were detected, whereas the three marine sites yielded 147 BINs, and the only brackish site had 75 BINs detected. Fifty-six BINs were shared between freshwater and marine sites, 46 between brackish and freshwater and 40 between brackish and marine sites (Figure 5, Table S4). If we consider the species ecology for each BIN, the brackish site included 44 freshwater taxa, 15 marine taxa and 3 organisms that can be found in both freshwater and marine environments. It should also be highlighted that although just a few marine taxa were detected in freshwater sites, as many as 53 freshwater BINs were detected in the three marine sites (Table S4). Even though we sampled water, 133 BINs detected were terrestrial taxa, mainly belonging to arthropods and annelids (Table S5). Among them, 125 were detected in freshwater sites, 11 were present in marine locations and 7 in the brackish site (Table S5). Thus, by subtracting these terrestrial taxa, the effective number of aquatic BINs detected in freshwater sites was 271, the number of aquatic BINs in marine sites was 136, and finally, 68 aquatic BINs were detected in the brackish site.

**Figure 5:**
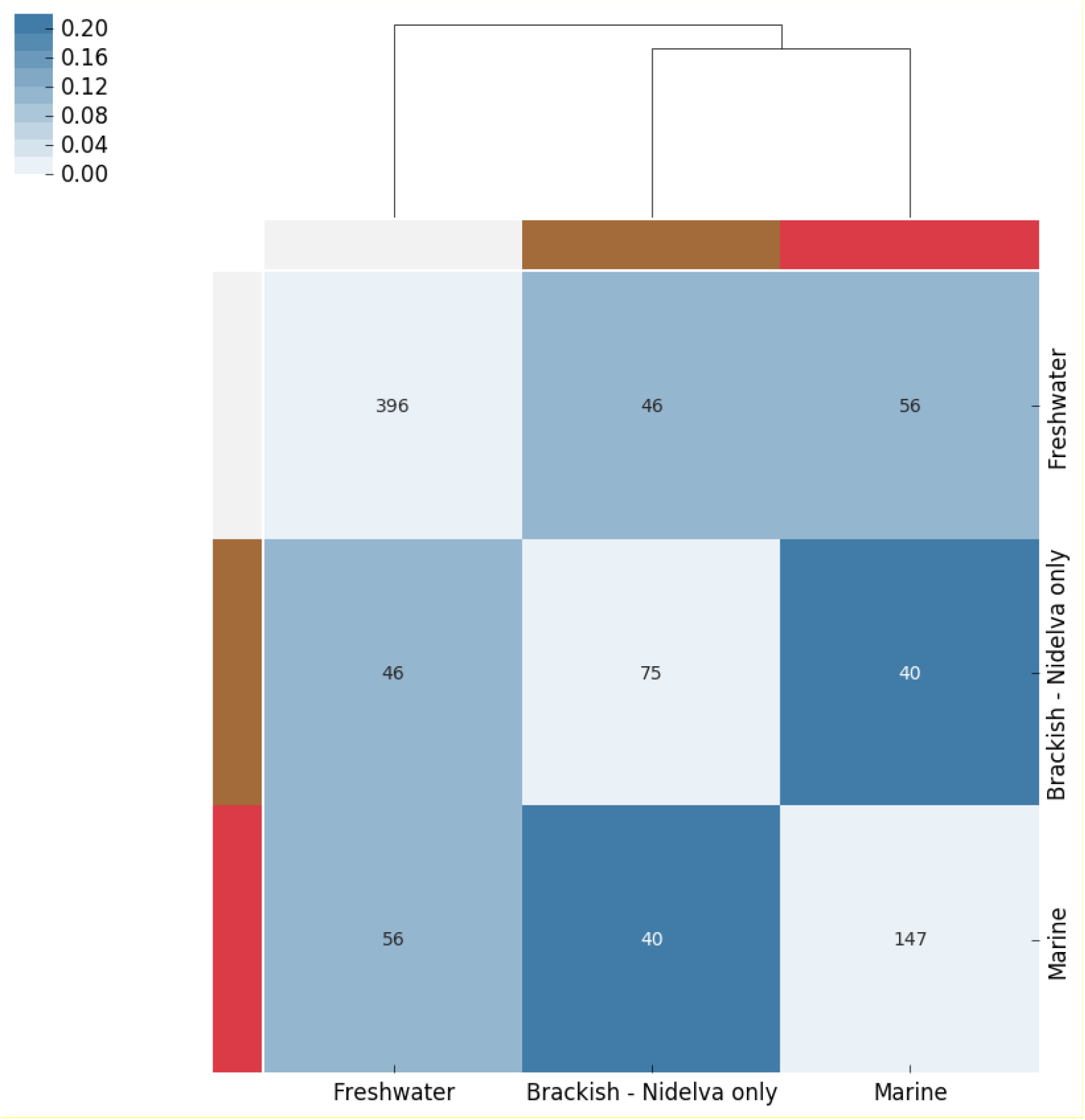
Heatmap generated in mBRAVE of Jaccard’s similarity index among habitat types. The darker the blue the greater the similarity is between sampled habitats. Clusters indicated with a dendrogram on top. Numbers between the habitat types correspond to the number of Barcode Index Numbers (BINs) shared. Diagonal numbers reflect the total number of BINs for each habitat type.

### 3.4 Transportation of eDNA signal along and between water bodies

The number of BINs detected differed along the Nidelva river. The highest number was 75 BINs detected downstream, at Sjøfartsmuseet (TRD_15/16), whereas 48 BINs were detected in the middle section of the stream, at Fylkesmannsboligen (TRD_17/18), and 27 upstream, at Sluppen (TRD_20-21). The Jaccard similarity index indicated a higher similarity between the downstream and middle parts of the river, compared to the upper part of Nidelva. Out of all BINs detected upstream, nearly half of them were shared between the middle part of the stream and the downstream Nidelva (13 and 14 BINs, respectively, Figure 6a) and the middle part shared 22 BINs with the downstream site. Among the detected BINs, seven BINs from downstream, seven BINs from Nidelva’s middle part and three BINs from upstream were terrestrial taxa and thus, trimming the observed diversity effectively to 68, 41 and 24 aquatic BINs, respectively. From the terrestrial habitat, only human DNA was shared among the three sites. Two more BINs (corresponding to the cow, *Bos taurus* Linnaeus and the red worm, *Eiseniella tetraedra* (Savigny)) were present in both the middle stream and downstream. The rest of the terrestrial taxa were found only in one of the three parts of the river and therefore, the terrestrial organisms did not have a profound effect on the similarities and the number of shared BINs reported above.

**Figure 6:**
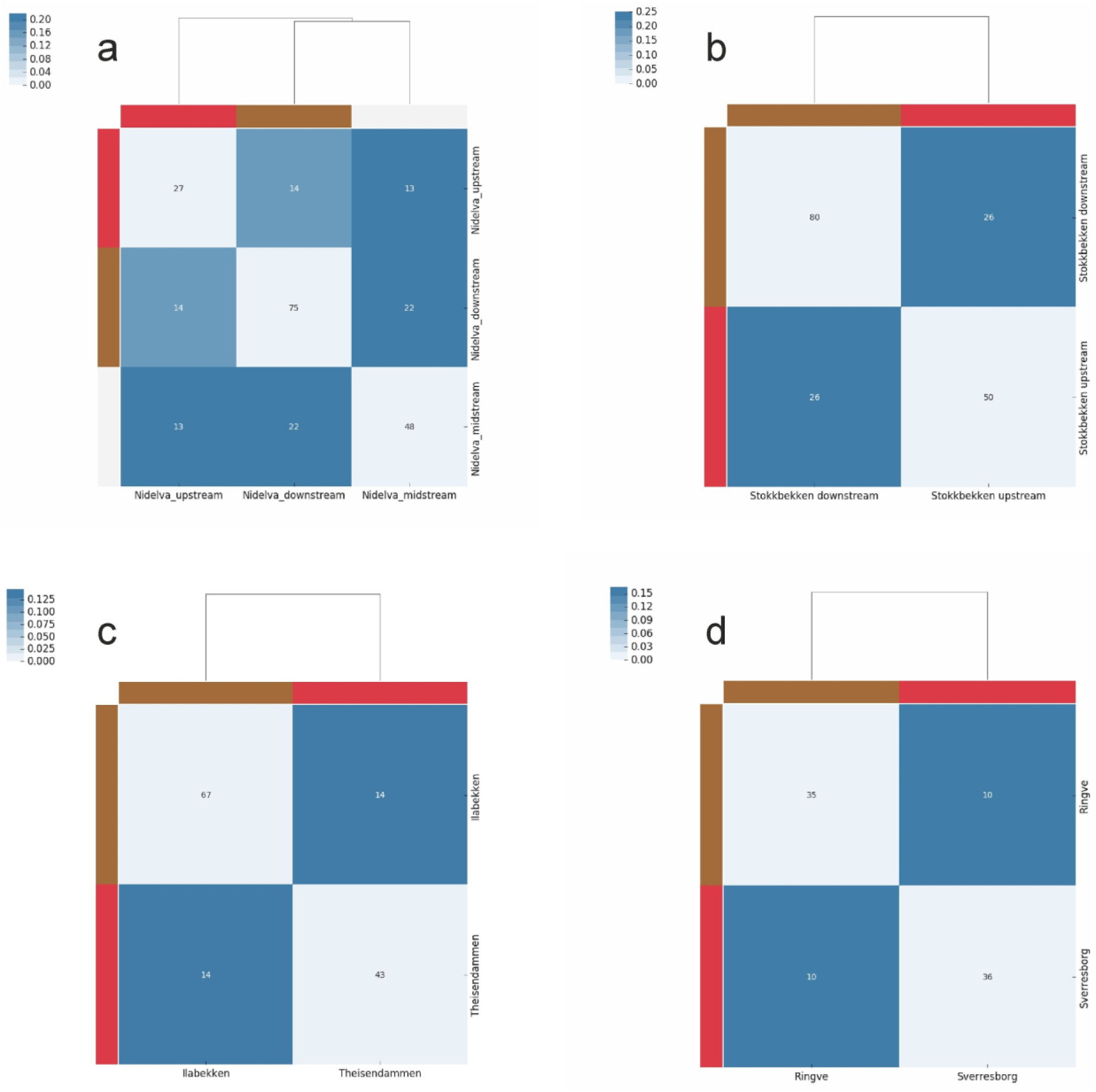
Heatmaps generated in mBRAVE of Jaccard’s similarity index among sites in selected water bodies. The darker the blue the greater the similarity is between sampled sites. Clusters indicated with a dendrogram on top. Numbers correspond to the number of Barcode Index Numbers (BINs) shared between respective sites. Diagonal numbers reflect the total number of BINs for each respective site.

Regarding the stream Stokkbekken (TRD_24-27), 50 BINs were detected downstream and 80 BINs in the upstream site. The Jaccard similarity value reached 0.25, with 26 BINs shared among the sites (Figure 6b). Among the detected taxa, 16 BINs from downstream and 15 BINs from upstream were terrestrial species and thus, the observed aquatic diversity was reduced to 34 BINs downstream and 65 BINs upstream. Seven terrestrial taxa were shared between the sites, including DNA from: human, four earthworms of the family Lumbricidae, one moth, often present at riverbanks, and one duck species. Therefore, both riverine sites shared 22 aquatic organisms. Theisendammen (TRD_3/4) contained 47 BINs, whereas in the stream Ilabekken (TRD_7/8) 67 BINs were detected. The Jaccard similarity value was 0.15, with 14 detected BINs shared between the sites (Figure 6c). Six BINs from Lake Theisendammen and 16 BINs from the stream Ilabekken were terrestrial taxa, therefore, the observed aquatic diversity was 41 and 51 BINs, respectively. Only one BIN, belonging to order Collembola, was shared between these sites. No common roach was detected.

The diversity in the two ponds at Sverresborg Museum (TRD_5/6) and Ringve Botanical Garden (TRD_13/14) was similar, with 36 and 35 BINs detected. Among those, 10 were shared between the ponds (Figure 6d), including DNA from five arthropods, two annelids, two proteobacteria and human, reaching the Jaccard similarity index value of 0.16. Within those BINs, terrestrial taxa were represented only by human DNA in the pond at Sverresborg and 2 BINs in the pond at Ringve Botanical Garden, including DNA from human and one duck species. Common newt (*Lissotriton vulgaris*) was detected in the pond at Sverresborg Museum.

In the single pond Madsjøen (TRD_22/23) 101 BINs were detected, whereas 130 BINs were present in its underground outflow Leangenbekken (TRD_28/29). The Jaccard similarity value reached 0.25, with 48 BINs shared among sites. Among the detected BINs, 58 BINs from pond Madsjøen and 63 BINs from Leangenbekken were terrestrial taxa, thus the observed diversity of aquatic BINs was 43 and 67 BINs, respectively. A total of 20 terrestrial taxa were detected from both sites. Great-crested newt was not detected in the pond Madsjøen.

## 4 Discussion

In this study, we demonstrate how aquatic eDNA can be sampled and sequenced to rapidly infer both terrestrial and aquatic diversity in the city of Trondheim, Norway. From just 30 litres of water sampled by two people in six hours across the city from different habitats, and sequenced within a short time, 435 species spanning 18 phyla could be detected from the total number of sequences obtained. Few detailed analyses of BioBlitz have been performed, but if we consider how many species on average were observed (an estimated 431.2 species per event) in the United Kingdom between 2006 and 2013 (Postles & Bartlett, 2018), then our biodiversity observations are equivalent. Additionally, because the data collected here was instantly digitized in the form of DNA sequences and available through mBRAVE, as a team of people from 14 different countries, we could engage in learning about the biodiversity in the city of Trondheim from all over the world. Although participants in BioBlitz events certainly enjoy observing biodiversity itself, there is educational value and excitement in detecting biodiversity that you cannot immediately observe when visiting a locality, as well as learning about biodiversity of places you could never visit.

As our results suggest, using an eDNA rapid biodiversity assessment in urban settings allows many basic questions about biodiversity to be assessed in and across a landscape. Patterns in overall diversity can be compared allowing educators to use findings to inform people and students about biodiversity. For example, the distribution of detected diversity matches that to known biodiversity patterns between land, marine, and freshwater. It holds that about 80% of species diversity is on land, 15% is in the oceans and the remaining 5% is in freshwater (Grosberg et al., 2012). Interestingly, even though we sampled water we still had a high proportion of DNA from terrestrial species including vertebrates such as deer and badgers. Recent studies focused on detection of terrestrial vertebrates from water samples show that it is a powerful way to non-invasively detect mobile and elusive vertebrate species (Harper et al., 2019). However, while vertebrate species detections are exciting, most detections were dominated by terrestrial beetles and annelids. The finding of Diptera as the dominant group of arthropods can be ascribed the high diversity of the midge family Chironomidae where most species have aquatic life stages.

Many improvements to sampling could lead to greater species detection. For example, of the sites sampled, 11 were freshwater, three were considered marine and one was classified as brackish, yet the number of detected species reflect the expected patterns of diversity, indicating that had a greater sampling effort occurred in marine and brackish sites, a greater amount of diversity would be detected as expected from the 15 % known diversity in these habitats. Additionally, our sampling was performed at one point in the season (mid-May), with low activity and abundance of some invertebrate groups at this high latitude (63° North). Thus, a greater number of taxa would likely be recorded had our sampling been done at several occasions throughout the seasons, due to differences in the life histories of present taxa. We also acknowledge the observation that we had a very low overlap in diversity among replicates (either biological or technical). Thus, to estimate total diversity even at a site, a greater number of samples from each site or greater sequencing depth could have been performed to increase the likelihood of detections.

In addition to the terrestrial species DNA being sampled from water, there was also evidence of DNA moving across the landscape. Examples include DNA from the lentic opossum shrimp (*Mysis relicta* Lovén) at all three sites in river Nidelva, DNA from the lotic mayfly *Baetis rhodani* Pictet in the lake Theisendammen as well as lotic caddisfly *Hydropsyche siltalai* Doehler in the Havstein pond, and about 20 non-biting midge taxa in the marine sites. Even though the transport of flying insects’ DNA between the sites can be easily explained, the presence of lentic mysid in the Nidelva river is surprising. Most likely, it can be associated with passive transport via man-made pipes distributing water from the Lake Selbusjøen, where it is known to occur, to the nearby power plant, which then potentially releases the water to upstream Nidelva.

Movement of DNA by the movement of water allows eDNA to measure biodiversity for scales larger than a single sampling site (Deiner et al., 2016). In this study, the sites sampled along the Nidelva river showed higher diversity downstream and a greater number of species were shared the further downstream the site was on the river, suggesting eDNA accumulation. However, one should bear in mind that transport is not necessarily confirmed with these patterns because the species detected have ecologies that would predict their occurrence in both sites along the river and can often be found present at higher densities downstream due to invertebrate drift (Grossman et al., 2010). The terrestrial and freshwater species DNA found in marine sites are more convincing examples of DNA being moved across the landscape. This inherent trait of eDNA moving away from its source is exciting when large areas need documenting and useful for BioBlitz events. But can also be troublesome when specific locations for a species are desired (e.g., as might be the case for an invasive species).

Among the three species of concern (two newts and the common roach), we detected the common newt (*Lissotriton vulgaris*) in the pond at Sverresborg cultural museum. The pond itself was restored in 2013-14 as part of the open air exhibition, and care was taken to preserve the known newt population (https://sverresborg.no/beskrivelse-av-dammen-og-parken). It is nice to see that this species presence in the pond could be confirmed by analysis of eDNA. Other vertebrates (fish, amphibians, birds and mammals) were detected at expected locations (Table S2), but it is noteworthy that DNA from common frog (*Rana temporaria* Linnaeus) and pig (*Sus scrofa* Linnaeus) were only found at the marine site at Korsvika. As there are no obvious breeding sites for frogs near the sampling site, the DNA of both species is likely of terrestrial origin. As Korsvika is a popular site for bathing and barbecuing, we suspect that pig DNA originates from human activities in the area. Alternatively, reagents used in PCR can also introduce animal DNA contamination is known from ancient DNA studies (Leonard et al., 2007), so we can not entirely rule this out as an explanation.

Reliable reference databases are needed to accurately assign a taxonomic name with an observed sequence (Weigand et al., 2019). Building such databases is underway, but they still need improvement. For example, we observed a significant level of taxonomic incongruences in the BIN assignment of sequences in our dataset. The most frequent misidentification observed was BIN discordance. BIN discordance is when one BIN is assigned to two or more species, and this was the case for more than 15% of all identified BINs. These discordant observations match a common pattern observed and discussed in numerous studies in various taxonomic groups (e.g., Bridge et al., 2003; Vilgalys, 2003;

Nilsson et al., 2006). Correct identification of the specimens present in the reference libraries relies heavily on the taxonomic expertise of the researchers, especially for closely-related, morphologically similar taxa. Since the results obtained in this study depend solely on the BOLD database, a certain level of misidentified sequences reflects the BIN discordance observed (Meiklejohn et al., 2019). We have also identified cases where one species comprised more than one BIN. Besides this being the result of misidentification cases, those might also reflect a certain level of hidden, yet undescribed cryptic diversity, which also confirms a global phenomenon observed across the tree of life (Fišer et al., 2018). For example, we have observed a high level of intraspecific diversity in a cosmopolitan and widespread worm species, *Tubifex tubifex* (Müller), identifying multiple BINs assigned to this taxon (Table S3). This finding is well supported by previous studies identifying substantial cryptic diversity in this species (e.g., Beauchamp et al., 2001), confirming that rapid eDNA surveys might also serve as a starting point for future, more detailed studies of local population level genetic diversity (Sigsgaard et al., 2017). However, one should be cautious about applying strict delimitation thresholds, which are far from universal to all taxonomic groups (e.g. Lin et al., 2015; Kvist, 2016).

In this survey we highlight mBRAVE, an open access computing platform with an interactive interface that performs metabarcoding bioinformatic processing. This platform allows parallel sequence identification for the COI barcoding region by comparison to the Barcode of Life Database for millions of sequences simultaneously. The significance of the mBRAVE platform is that it allows archiving, sharing and exploration of complex genetic data through visualizations and tools that require no computer science background and can be conducted online. It is noted that further diversity analyses are possible outside the mBRAVE environment by downloading the data and using it elsewhere, but no such external analyses were performed for the purpose of this study.

## 5 Conclusions

Our goal was to demonstrate the use of DNA-based detection methods for urban biodiversity surveys and suggest ways for environmental educators in how they can run their own rapid biodiversity surveys using eDNA. We recommend using our study as an example and suggest improvements for how to implement the field and lab protocols. Although, the latter can be done in collaboration with a local university or company who perform such analysis. While these studies can be carried out in the absence of direct interaction with taxonomic experts due to the digitisation of current knowledge in the Barcode of Life Database through DNA barcoding, it is still highly encouraged to collaborate with them. Even though these DNA-based methods are exciting, they are heavily reliant on taxonomic identifications provided by a reliable reference database. While much work remains to perfect our abilities for DNA-based detection of life, surveys of this type are useful and exciting explorations for building an engaged public interested in reversing the loss of life from our planet.

## Data Accessibility

The data is publicly available within the mBRAVE environment (Project Code: MBR-WRKMISEQPE), and in the European Nucleotide Archive (ENA) sequence read archive (PRJEB37589).

## Conflicts of interest

None declared

## Acknowledgements

We thank the NTNU University Museum for hosting the mBRAVE scientific workshop in conjunction with the 8th International Barcode of Life Conference. We are grateful for the sequencing services provided by the Helse Sør-Øst Genomics Core Facility at Oslo University Hospital. VWR, Illumina, Wiley for financial support, Leonardo A. Meza-Zepeda for squeezing in our library for sequencing and Sujeevan Ratnasingham, Megan Milton, Tony Kuo for providing the workshop. We also thank Marc Daverdin, NTNU University Museum for creating the map in Fig. 1.

## Authors’ contributions

Sampling, laboratory work and study design: MM, TE

Data analyses: KH, MM, MC, LS, DM, VI, DAL, SL, FCJ, ZCZ, ISM, AD, LT, KM, DH, GK, XL, BP, TE, KD

Supervision: KH, KD

Manuscript preparation: all authors edited and commented on manuscript

---

Table S1: Collection sites of sampled eDNA

Table S2: Taxonomic assignment and habitat association of retrieved Barcode Index Numbers (BINs) per location.

Table S3: Taxonomic incongruences of retrieved Barcode Index Numbers (BINs) in the BOLD reference database.

Table S4: Shared Barcode Index Numbers (BINs) between the sites representing different habitats.

Table S5: Terrestrial Barcode Index Numbers (BINs) retrieved per location.

Table S1.1: Indexing scheme used for this study.

Table S1.2: Results of quality filtering sequences for an example run of TRD01_1_Dam_Havstein_golfbane.

Table S2.1: The impact of singleton and doubleton removal on total number of Barcode Index Numbers (BINs), their taxonomic rank assignment and the percentages of shared BINs among the biological and technical samples for a site when the BIN was shared by at least two replicates within a sampling site.

Table S2.2 Barcode Index Numbers (BINs) present in the blank samples with number of sequences in each replicate.

## Notes

### Competing Interest Statement

The authors have declared no competing interest.

## References

Ashfaq M., Sabir J.S.M., El-Ansary H.O., Perez K., Levesque-Beaudin V., Khan A.M., Rasool A., Gallant C., Addesi J., Hebert P.D.N. (2018). Insect diversity in the Saharo-Arabian region: Revealing a little-studied fauna by DNA barcoding. PLoS One, 13, 1–16, e0199965, DOI: 10.1371/journal.pone.0199965

Beauchamp K.A., Kathman R.D., McDowell T.S., Hedrick R.P. (2001). Molecular phylogeny of tubificid oligochaetes with special emphasis on *Tubifex tubifex* (Tubificidae). Molecular Phylogenetics and Evolution, 19(2), 216–224.

Bridge P. D., Roberts P. J., Spooner B. M., Panchal G. (2003). On the unreliability of published DNA sequences. New Phytologist, 160(1), 43–48.

Cristescu M.E., Hebert P.D. (2018). Uses and misuses of environmental DNA in biodiversity science and conservation. Annual Review of Ecology, Evolution, and Systematics, 49, 209–230.

D’Souza M.L., Hebert P.D.N. (2018). Stable baselines of temporal turnover underlie high beta diversity in tropical arthropod communities. Molecular Ecology, 27, 2447–2460.

Deiner K., Fronhofer E., Mächler E., Walser J.C., Altermatt F. (2016). Environmental DNA reveals that rivers are conveyer belts of biodiversity information. Nature Communications, 7, 12544.

Deiner K., Bik H.M., Mächler E., Seymour M., Lacoursière-Roussel A., Altermatt F., … & Pfrender M.E. (2017). Environmental DNA metabarcoding: Transforming how we survey animal and plant communities. Molecular Ecology, 26(21), 5872–5895.

de Jong Y., Verbeek M., Michelsen V., de Place Bjørn P., Los W., Steeman F., … & Hagedorn G. (2014). Fauna Europaea-all European animal species on the web. Biodiversity Data Journal, 2: e4034. doi: 10.3897/BDJ.2.e4034.

Elbrecht, V., Leese F. (2017). Validation and development of COI metabarcoding primers for freshwater macroinvertebrate bioassessment. Frontiers in Environmental Science 10, https://doi.org/10.3389/fenvs.2017.00011.

Fišer C., Robinson C.T., Malard F. (2018). Cryptic species as a window into the paradigm shift of the species concept. Molecular Ecology, 27(3), 613–635.

Grosberg R.K., Vermeij, G.J., Wainwright, P.C. (2012). Biodiversity in water and on land. Current Biology, 22(21), R900–R903. doi:10.1016/j.cub.2012.09.050

Grossman G.D., Ratajczak Jr. R.E., Farr M.D., Wagner C.M., Petty J.T. (2010). Why There Are Fewer Fish Upstream? American Fisheries Society Symposium, 73, 63–81.

Harper L.R., Handley L.L., Carpenter A.I., Ghazali M., Di Muri C., Macgregor C.J., Logan T.W., Law A., Breithaupt T., Read D.S., McDevitt A.D. (2019). Environmental DNA (eDNA) metabarcoding of pond water as a tool to survey conservation and management priority mammals. Biological Conservation, 238, 108225.

Hebert P.D.N., Ratnasingham S., Zakharov E.V., Telfer A.C., Levesque-Beaudin V., Milton M.A., Pedersen S., Jannetta P., deWaard J.R. (2016). Counting animal species with DNA barcodes: Canadian insects. Philosophical Transactions of the Royal Society B: Biological Sciences, 371(1702), 10.

Horton T., Gofas S., Kroh A., Poore G.C., Read G., Rosenberg G., … & Costello M.J. (2017). Improving nomenclatural consistency: a decade of experience in the World Register of Marine Species. European Journal of Taxonomy, 389, 1–24.

IPBES (2019). Global assessment report on biodiversity and ecosystem services of the Intergovernmental Science-Policy Platform on Biodiversity and Ecosystem Services. E. S. Brondizio, J. Settele, S. Díaz, and H. T. Ngo (editors). IPBES secretariat, Bonn, Germany

Kvist S. (2016). Does a global DNA barcoding gap exist in Annelida?. Mitochondrial DNA Part A, 27(3), 2241–2252.

Leonard J.A., Shanks O., Hofreiter M., Kreuz E., Hodges L., Ream W., … & Fleischer R. C. (2007). Animal DNA in PCR reagents plagues ancient DNA research. Journal of Archaeological Science, 34(9), 1361–1366.

Lin X., Stur E., Ekrem T. (2015). Exploring Genetic Divergence in a Species-Rich Insect Genus Using 2790 DNA Barcodes. PloS one 10(9), e0138993, DOI: 10.1371/journal.pone.0138993

Meiklejohn K.A., Damaso N., Robertson J.M. (2019). Assessment of BOLD and GenBank–Their accuracy and reliability for the identification of biological materials. PloS one, 14(6), e0217084, DOI: 10.1371/journal.pone.0217084

Nilsson R.H., Ryberg M., Kristiansson E., Abarenkov K., Larsson K.H., Kõljalg U. (2006). Taxonomic reliability of DNA sequences in public sequence databases: a fungal perspective. PloS one, 1(1), : e59, DOI: 10.1371/journal.pone.0000059

Postles M., Bartlett M. (2018). The rise of BioBlitz: Evaluating a popular event format for public engagement and wildlife recording in the United Kingdom, Applied Environmental Education & Communication, 17:4, 365-379, DOI: 10.1080/1533015X.2018.1427010

Ratnasingham S., Hebert P.D. (2007). BOLD: The Barcode of Life Data System (http://www.barcodinglife.org). Molecular Ecology Notes, 7(3), 355–364.

Ratnasingham S., Hebert, P.D. (2013). A DNA-based registry for all animal species: the Barcode Index Number (BIN) system. PloS one, 8(7), e66213. DOI: 10.1371/journal.pone.0066213

Ruch D.G., Karns D.R., McMurray P., Moore-Palm J., Murphy W., Namestnik S.A., Roth K. (2010). Results of the Loblolly Marsh Wetland Preserve BioBlitz, Jay County, Indiana. Proceedings of the Indiana Academy of Science, 119(1), 1–3.

Sigsgaard E.E., Nielsen I.B., Bach S.S., Lorenzen E.D., Robinson D.P., Knudsen S.W., Pedersen M.W., Al Jaidah M., Orlando L., Willerslev E., Møller P.R. (2017). Population characteristics of a large whale shark aggregation inferred from seawater environmental DNA. Nature ecology & evolution, 1(1), 0004.

Thomsen P.F., Sigsgaard E.E. (2019). Environmental DNA metabarcoding of wild flowers reveals diverse communities of terrestrial arthropods. Ecology and Evolution, 9(4), 1665–1679.

Tilseth E. (2008). Mapping of newt localities in the municipality of Trondheim 2007-2008. City of Trondheim, Department of Environment Report TM 2008/06, 237p.

Vilgalys R. (2003). Taxonomic misidentification in public DNA databases. New Phytologist, 160(1), 4–5.

Weigand H., Beermann A.J., Čiampor F., Costa F.O., Csabai Z., Duarte S., … & Strand M. (2019). DNA barcode reference libraries for the monitoring of aquatic biota in Europe: Gap-analysis and recommendations for future work. Science of the Total Environment, 678, 499–524.

