## Supplementary File 1 for "A rapid urban biodiversity blitz using aquatic environmental DNA"

Supplementary File 1. Detailed description of pipeline parameter settings used in the study.

S1.1 Merging and Trimming

After import of the raw fastq files into the mBRAVE environment. We performed the options needed for merging and trimming Illumina instrument produced sequence data. To merge the forward and reverse reads, we used an overlap of at least 20 base pairs to be certain that they are merged at homologous positions into a single sequence. Additionally, reads that were expected to merge were excluded if they had more than 5 nucleotide substitutions.

Given that raw reads often have indices and primers still attached, the sequences were trimmed by removing the front and tail end of the merged reads. Given that the indices (Table S1.1) have already been trimmed by the software used to dereplicate the libraries on the Illumina MiSeq, here we trimmed 20 bp of the front and the end of each sequence, representing the primers that were used for this study. We set up the trim length parameter to a total length of sequences down to 500bp, although the primers used target a region of around between 410 to 430 bp (~420 bp for most species). We did not use the primer masking function.

Table S1.1 Indexing scheme used for this study.

| **Locality** | **Replicate** | **I7 index** | **I5 index** |
| --- | --- | --- | --- |
| pond Havstein golf course | TRD1_1 | ACTCGCTA | TTATGCGA |
|  | TRD1_2 | CGGAGCCT | TCGACTAG |
|  | TRD1_3 | ACTGAGCG | GCGTAAGA |
|  | TRD2_1 | GGAGCTAC | TTATGCGA |
|  | TRD2_2 | GCGTAGTA | TCGACTAG |
|  | TRD2_3 | TACGCTGC | CCTAGAGT |
| Theisendammen | TRD3_1 | GCGTAGTA | CTATTAAG |
|  | TRD3_2 | TACGCTGC | TTCTAGCT |
|  | TRD3_3 | ACTGAGCG | TTATGCGA |
|  | TRD4_1 | ATGCGCAG | TTCTAGCT |
|  | TRD4_2 | ATGCGCAG | GAGCCTTA |
|  | TRD4_3 | ACTGAGCG | TTCTAGCT |
| pond Sverresborg museum | TRD5_1 | ACTCGCTA | CTATTAAG |
|  | TRD5_2 | CCTAAGAC | TTCTAGCT |
|  | TRD5_3 | TCGACGTC | GCGTAAGA |
|  | TRD6_1 | ACTGAGCG | TCGACTAG |
|  | TRD6_2 | CCTAAGAC | TTATGCGA |
|  | TRD6_3 | TCGACGTC | CCTAGAGT |
| Ilabekken at Bleikvollen | TRD7_1 | CGGAGCCT | CCTAGAGT |
|  | TRD7_2 | TAGCGCTC | TCGACTAG |
|  | TRD7_3 | TAGCGCTC | CCTAGAGT |
|  | TRD8_1 | ACTCGCTA | TTCTAGCT |
|  | TRD8_2 | TGCAGCTA | TCGACTAG |
|  | TRD8_3 | TGCAGCTA | AAGGCTAT |
| Pirsenteret | TRD9_1 | ACTCGCTA | TCGACTAG |
|  | TRD9_2 | CGGAGCCT | AAGGCTAT |
|  | TRD9_3 | TGCAGCTA | GCGTAAGA |
|  | TRD10_1 | TAGCGCTC | AAGGCTAT |
|  | TRD10_2 | CCTAAGAC | GAGCCTTA |
|  | TRD10_3 | TCGACGTC | TCGACTAG |
| Korsvika | TRD11_1 | GGAGCTAC | GCGTAAGA |
|  | TRD11_2 | CGGAGCCT | GAGCCTTA |
|  | TRD11_3 | TACGCTGC | GAGCCTTA |
|  | TRD12_1 | GCGTAGTA | GCGTAAGA |
|  | TRD12_2 | TAGCGCTC | TTCTAGCT |
|  | TRD12_3 | CGATCAGT | TTCTAGCT |
| Ringve Botanical Garden | TRD13_1 | CGGAGCCT | TTATGCGA |
|  | TRD13_2 | ATGCGCAG | AAGGCTAT |
|  | TRD13_3 | ACTGAGCG | AAGGCTAT |
|  | TRD14_1 | TACGCTGC | AAGGCTAT |
|  | TRD14_2 | CCTAAGAC | GCGTAAGA |
|  | TRD14_3 | TCGACGTC | CTATTAAG |
| Nidelva at sjøfartsmuseet | TRD15_1 | TACGCTGC | TCGACTAG |
|  | TRD15_2 | TAGCGCTC | TTATGCGA |
|  | TRD15_3 | TGCAGCTA | CCTAGAGT |
|  | TRD16_1 | ACTCGCTA | AAGGCTAT |
|  | TRD16_2 | CGATCAGT | TCGACTAG |
|  | TRD16_3 | CGATCAGT | CTATTAAG |
| Nidelva at fylkesmannsboligen | TRD17_1 | GGAGCTAC | GAGCCTTA |
|  | TRD17_2 | TACGCTGC | TTATGCGA |
|  | TRD17_3 | TAGCGCTC | GCGTAAGA |
|  | TRD18_1 | GGAGCTAC | CTATTAAG |
|  | TRD18_2 | GCGTAGTA | CCTAGAGT |
|  | TRD18_3 | CGATCAGT | GCGTAAGA |
| blank sample | TRD19_1 | ACTCGCTA | GCGTAAGA |
|  | TRD19_2 | CGATCAGT | GAGCCTTA |
|  | TRD19_3 | TGCAGCTA | GAGCCTTA |
| Nidelva at Sluppen | TRD20_1 | GGAGCTAC | CCTAGAGT |
|  | TRD20_2 | GCGTAGTA | TTCTAGCT |
|  | TRD20_3 | GCGTAGTA | TTATGCGA |
|  | TRD21_1 | TAGCGCTC | CTATTAAG |
|  | TRD21_2 | ACTGAGCG | GAGCCTTA |
|  | TRD21_3 | TCGACGTC | TTATGCGA |
| Madsjøen at IKEA | TRD22_1 | GCGTAGTA | GAGCCTTA |
|  | TRD22_2 | TACGCTGC | CTATTAAG |
|  | TRD22_3 | ATGCGCAG | CCTAGAGT |
|  | TRD23_1 | GGAGCTAC | TTCTAGCT |
|  | TRD23_2 | ATGCGCAG | TCGACTAG |
|  | TRD23_3 | ACTGAGCG | CCTAGAGT |
| Stokkbekken upstream | TRD24_1 | ATGCGCAG | GCGTAAGA |
|  | TRD24_2 | CCTAAGAC | TCGACTAG |
|  | TRD24_3 | TGCAGCTA | TTATGCGA |
|  | TRD25_1 | CGGAGCCT | GCGTAAGA |
|  | TRD25_2 | CCTAAGAC | CTATTAAG |
|  | TRD25_3 | CGATCAGT | CCTAGAGT |
| Stokkbekken downstream | TRD26_1 | GCGTAGTA | AAGGCTAT |
|  | TRD26_2 | CGATCAGT | TTATGCGA |
|  | TRD26_3 | TGCAGCTA | TTCTAGCT |
|  | TRD27_1 | ACTCGCTA | CCTAGAGT |
|  | TRD27_2 | CGGAGCCT | TTCTAGCT |
|  | TRD27_3 | ATGCGCAG | CTATTAAG |
| Leangenbekken | TRD28_1 | GGAGCTAC | AAGGCTAT |
|  | TRD28_2 | TAGCGCTC | GAGCCTTA |
|  | TRD28_3 | CCTAAGAC | AAGGCTAT |
|  | TRD29_1 | ACTCGCTA | GAGCCTTA |
|  | TRD29_2 | ATGCGCAG | TTATGCGA |
|  | TRD29_3 | TCGACGTC | TTCTAGCT |
| Østmarkneset at Ladekaia | TRD30_1 | GGAGCTAC | TCGACTAG |
|  | TRD30_2 | TCGACGTC | AAGGCTAT |
|  | TRD30_3 | TCGACGTC | GAGCCTTA |
|  | TRD31_1 | ACTGAGCG | CTATTAAG |
|  | TRD31_2 | CGATCAGT | AAGGCTAT |
|  | TRD31_3 | TGCAGCTA | CTATTAAG |
| blank sample | TRD32_1 | CGGAGCCT | CTATTAAG |
|  | TRD32_2 | TACGCTGC | GCGTAAGA |
|  | TRD32_3 | CCTAAGAC | CCTAGAGT |

S1.2 Filtering

The filtering of sequences is to prevent inclusion of artificial reads due to technical errors. During sequencing, each nucleotide in every sequence is evaluated and given a quality value (QV). This quality value reflects the probability of that nucleotide being the correct nucleotide in that position of the sequence. QV is used in the formula QV = -10log(*Pe*), where Pe is the probability of error. For example, a QV of 20 then means that there is a 1% chance that the sequencer reads a wrong nucleotide for the sequence in each position (1 in 100 nucleotides are wrong, given that they all have QV=20). Thus, we chose a minimum average 20 QV value, and allowed for a max of 2% of the nucleotides to be below this threshold. Additionally, we allowed max 1% of the nucleotides to be below QV 10. This filtering threshold removed about 30% of all sequences, changing the mean length of obtained sequences to 406bp, compared to the initial 360bp (Table S1.2).

Table S1.2 Results of quality filtering sequences for an example run of TRD01_1_Dam_Havstein_golfbane.

|  | Reads | Mean length | Mean QV | Mean GC Composition |
| --- | --- | --- | --- | --- |
| Pre-filtering | 208639 | 360.17bp | 39.07 | 42.77% |
| Post-filtering | 142600 | 406.31bp | 38.59 | 39.23% |

S1.3 OTU clustering and BIN association

We clustered Operational Taxonomic Units (OTUs) using a sequence similarity threshold of 2.5%, without using any pre-clustering threshold. This means that 2.5 % intraspecific variability was allowed within each OTU. For the taxonomic assignment, represented here by BIN association, we used a 2% similarity threshold, meaning an OTU must match a reference sequence with at least 98% of its nucleotides to be assigned to a taxonomic name from the associated reference sequence. The other settings describe the minimum number of sequences of that specific OTU required to be detected and included as a BIN for a given sample. If an OTU does not match a reference sequence within the 2% similarity criterion, mBRAVE analysis checks if there are any matches when the threshold is lower as follows:

- Species Similarity is based on the ID Distance threshold (2.0%).
- Genus Similarity is based on ID Distance threshold plus 1%.
- Subfamily Similarity is based on ID Distance threshold plus 2%.
- Family Similarity is based on ID Distance threshold plus 3%.
- Order Similarity is based on ID Distance threshold plus 5%.
- Class Similarity is based on ID Distance threshold plus 8%.
- Phylum Similarity is based on ID Distance threshold plus 10%.

The final part of our analysis involved the taxonomic assignment of the produced OTUs against reference libraries. The mBRAVE workflow runs all delimited OTUs against BOLD reference libraries derived custom datasets and runs independent identification algorithms against project libraries as described in the main text.
