## Supplementary File 2 for "A rapid urban biodiversity blitz using aquatic environmental DNA"

Supplemental File 2: Detailed analysis of shared Barcode Index Numbers (BINs) and sequence representation and discordance of taxonomic information associated with BINs.

S2.1 Impact of singleton/doubleton removal

When using more strict conditions and considering a particular BIN to be shared if it had been recovered in at least two replicates of either of the two samples from the same locality, then on average BINs were shared at 32% among replicates of particular sampling site, with the similarity ranging from 12% to 50% (Table S2.1).

Table S2.1: The impact of singleton and doubleton removal on total number of Barcode Index Numbers (BINs), their taxonomic rank assignment and the percentages of shared BINs among the biological and technical samples for a site when the BIN was shared by at least two replicates within a sampling site.

| **Removal** | **BINs** | **SP** | **G** | **F** | **O** | **C** | **P** | **Sequences** | **Av. %**  **shared BINs** | **Max % shared BINs** | **Min % shared BINs** |
| --- | --- | --- | --- | --- | --- | --- | --- | --- | --- | --- | --- |
| **None** | 501 | 435 | 324 | 184 | 90 | 39 | 18 | 552277 | 32.14 | 50.98 | 12.00 |
| **Singleton** | 490 | 425 | 317 | 179 | 87 | 38 | 18 | 552097 | 32.50 | 55.56 | 9.09 |
| **Doubleton** | 469 | 410 | 306 | 171 | 83 | 38 | 18 | 551947 | 33.14 | 53.33 | 5.00 |

The total number of detected BINs was different when singletons and doubletons were removed from the dataset. The overall number of BINs dropped down to 490 and 469, respectively, with decline observed also in the number of species (425 and 410) and the orders (87 and 83), whereas the number of phyla detected remained the same in both cases (Table S2.1). Interestingly, it mainly affected the BIN composition of the blank samples. When singletons were removed, only two BINs were retrieved from blank samples, being Alphaproteobacteria (TAX:701540) and Diptera (BOLD:ADT6228), and when the doubletons were excluded solely the Diptera BIN remained in the blanks (Table S2.2). Although the singleton and doubleton removal did not affect the average similarity between replicates within a site, it altered the ranges of values of shared BINs, by raising the maximum values (Table S2.1).

Regarding the similarity values within the groups indicated by Jaccard similarity index, they were not significantly altered by the singleton and doubleton removal. However, the index value did change between groups, making them more distinguishable from each other when singletons and doubletons were removed.

Table S2.2 Barcode Index Numbers (BINs) present in the blank samples with number of sequences in each replicate.

| **Phylum** | **Order** | **BIN URI** | **Samples** | | | | | |
| --- | --- | --- | --- | --- | --- | --- | --- | --- |
|  |  |  | 19-1 | 19-2 | 19-3 | 32-1 | 32-2 | 32-3 |
| Annelida | Haplotaxida | BOLD:AAA6233 | 0 | 0 | 0 | 1 | 0 | 0 |
| Annelida | Haplotaxida | BOLD:ACB6471 | 0 | 0 | 0 | 1 | 0 | 0 |
| Arthropoda | Diplostraca | BOLD:ACF5469 | 0 | 0 | 0 | 0 | 1 | 0 |
| Arthropoda | Coleoptera | BOLD:AAO3823 | 0 | 0 | 0 | 0 | 1 | 0 |
| Arthropoda | Coleoptera | BOLD:AAH2826 | 0 | 0 | 0 | 0 | 0 | 1 |
| Arthropoda | Diptera | BOLD:ADT6228 | 0 | 0 | 203 | 142 | 9 | 0 |
| Arthropoda | Plecoptera | BOLD:ACY3863 | 0 | 0 | 0 | 1 | 0 | 0 |
| Proteobacteria |  | TAX:701540 | 1 | 2 | 0 | 0 | 0 | 0 |
| Rotifera | Ploima | BOLD:ACL8151 | 0 | 0 | 0 | 0 | 1 | 0 |

S2.2 BIN discordance and errors in taxonomic names

Interestingly, there were numerous cases of BIN discordance (all listed in Table S3). There were as many as 80 cases where one BIN is assigned to two or more distinct species, with most of them occurring in the arthropod BINs.

Moreover, we identified 21 species with more than one BIN reaching maximum up to seven distinct BINs within one recognized species. Most of those cases involved annelid species with notable cases in arthropods and rotifers as well.

We identified also two cases where likely misspelling led to the overrepresentation of species belonging to one BIN, namely BOLD:AAD4606 (‘*Podon leuckarti*’ and ‘*Podon leuckartii*’) and BOLD:AAK2653 (‘*Capnopsis schilleri*’ and ‘*Capnopsis chilleri*’).
