## Supplementary material for "A rapid urban biodiversity blitz using aquatic environmental DNA": Table S1

Table S1 Collection sites of sampled eDNA

| **Site Code** | **Date of collection** | **Locality** | **Latitude** | **Longitude** | **Habitat** |
| --- | --- | --- | --- | --- | --- |
| TRD_1/2 | 13.05.2019 | pond Havstein golf course, Trondheim | 63.408151 | 10.371569 | freshwater |
| TRD_3/4 | 13.05.2019 | Theisendammen, Trondheim | 63.422701 | 10.346773 | freshwater |
| TRD_5/6 | 13.05.2019 | pond Sverresborg museum, Trondheim | 63.417837 | 10.354383 | freshwater |
| TRD_7/8 | 13.05.2019 | Ilabekken at Bleikvollen, Trondheim | 63.42933 | 10.36359 | freshwater |
| TRD_9/10 | 13.05.2019 | Pirsenteret, Trondheim | 63.441681 | 10.401761 | marine |
| TRD_11/12 | 13.05.2019 | Korsvika, Trondheim | 63.450008 | 10.433459 | marine |
| TRD_13/14 | 13.05.2019 | Ringve Botanical Garden, Trondheim | 63.448969 | 10.453557 | freshwater |
| TRD_15/16 | 13.05.2019 | Nidelva at sjøfartsmuseet, Trondheim | 63.434292 | 10.406706 | brackish |
| TRD_17/18 | 13.05.2019 | Nidelva at fylkesmannsboligen, Trondheim | 63.428981 | 10.379226 | freshwater |
| TRD_19 | 13.05.2019 | blank sample |  |  |  |
| TRD_20/21 | 14.05.2019 | Nidelva at Sluppen, Trondheim | 63.395789 | 10.386931 | freshwater |
| TRD_22/23 | 14.05.2019 | Madsjøen at IKEA, Trondheim | 63.427133 | 10.475400 | freshwater |
| TRD_24/25 | 14.05.2019 | Stokkbekken upstream, Trondheim | 63.419893 | 10.490933 | freshwater |
| TRD_26/27 | 14.05.2019 | Stokkbekken downstream, Trondheim | 63.434519 | 10.499838 | freshwater |
| TRD_28/29 | 14.05.2019 | Leangenbekken, Trondheim | 63.438268 | 10.473162 | freshwater |
| TRD_30/31 | 14.05.2019 | Østmarkneset at Ladekaia, Trondheim | 63.457253 | 10.446165 | marine |
| TRD_32 | 14.05.2019 | blank sample |  |  |  |
